# In situ radiation damage tracking during scanning small-angle X-ray scattering

**DOI:** 10.64898/2026.10.20.760376

**Authors:** Boram Yu, Rita Mendes Da Silva, Mangalika Sinha, Ulrike Rölleke, Manfred Burghammer, Sarah Köster

## Abstract

Due to the high penetration power of X-rays, X-ray imaging techniques have been employed to image cellular architecture. Among these techniques, scanning small-angle X-ray scattering (SAXS) is a powerful method for investigating ordered structures within cells. With the recent advent of high-brilliance synchrotron sources, nano-focused X-rays and fast scanning modes, scanning SAXS can now provide structural information from hydrated cells in liquid environments in little time. However, the high photon flux results in an increased dose and pronounced radiation damage to the intracellular structures. Therefore, strategies are required to monitor and quantitatively analyze the extent of damage. Here, we present a multimodal approach that employs an on-axis, high-speed, brightfield microscope to monitor cellular damage during scanning. This microscope is integrated into the scanning SAXS setup at ID13 of ESRF and records images in synchrony with SAXS data acquisition. We are able to observe and quantitatively track changes in cellular morphology *in situ* and thereby obtain complementary information to that provided by scanning SAXS measurements. Our approach opens up the potential for a multifaceted investigation of radiation damage to biological cells and other sample types.

**Synopsis:** We present a versatile *operando* light microscopy method to track radiation damage in sensitive biological samples during X-ray imaging, including adapted data analysis concepts.

## 1 Introduction

Imaging is a key technique for investigating cellular architecture and function. Using X-rays for cellular imaging has several advantages, among them the short wavelength and high penetration power, which enables imaging with high spatial resolution of whole cells without sectioning. Additionally, X-ray imaging is label-free and does not require staining when preparing samples. Among the various X-ray imaging techniques, scanning small-angle X-ray scattering (SAXS) provides real-space images with moderate resolution as well as reciprocal-space information with nanoscale resolution. Thus, scanning SAXS has been used to study ordered structures within cells (Weinhausen *et al*., 2012; Poccia *et al*., 2014; Hémonnot *et al*., 2016; Bernhardt *et al*., 2017; Nicolas *et al*., 2019*b*). The recent advent of brighter X-rays in large X-ray facilities enables fast scanning and thereby high-throughput data acquisition in less time (Cassini *et al*., 2020; Yu *et al*., 2026).

A critical aspect for cellular imaging with X-rays is radiation damage. X-radiation damages cellular components, especially at high doses (Nicolas *et al*., 2019*a*; Yu *et al*., 2026). This consideration must be taken into account even more carefully when measuring cells in liquid environments. These environments provide close-to-physiological conditions for cells and establish an important step towards live-cell investigations. However, the SAXS signal of biological cells in aqueous environments is inherently weak and the radiation damage is more pronounced than for dried samples (Riley, 1994; Le Caër, 2011; Hémonnot & Köster, 2017) because of radiolysis. Therefore, it is important to optimize the measurement parameters, such as X-ray energy, scan step size and exposure time, in order to record high-quality data while minimizing the dose. To this end, a method to monitor *in situ* and quantify the extent of damage is required. Various methods have been proposed to observe and quantify damage, including tracking changes in chemical composition (Kosior *et al*., 2012; Lin *et al*., 2024), morphology (Gianoncelli *et al*., 2015; Strelnikova *et al*., 2017), or intracellular structure (Nicolas *et al*., 2019*a*; Yu *et al*., 2026) at different radiation doses or after repeated scans. However, these studies only monitor changes after each exposure, not during the process. Thus, these approaches do not support monitoring changes *in situ* and, thereby, cannot be used to control the damage during a measurement.

On-axis microscopes, *i*.*e*., integrated in the beamline for *in situ* use, have been presented for monitoring the sample condition during synchrotron X-ray measurements. These microscopes used a dichroic mirror (Madden *et al*., 2013), a right-angle mirror (van Silfhout, 2021; Liu *et al*., 2023), or a 45-degree mirror (Schneider *et al*., 2021) with a drilled hole to allow the X-rays to pass through. These microscopes are primarily designed to monitor the changes in sample state and to determine the correct timing for the measurements. Therefore, they are used as auxiliary tools rather than for collecting images. For fast scanning SAXS measurements with on-axis microscopy, in particular, the microscope requires a wide field of view to cover the entire scan area in a single view, as well as high-speed imaging techniques for rapid image storage and processing.

Here, we present a high-speed, on-axis, brightfield microscope integrated into a scanning SAXS setup at ID 13, ESRF (Grenoble, France), including the design and commissioning of the microscope and corresponding data analysis for monitoring and quantitatively tracking the radiation damage to the cells during the scanning process. We compare the changes observed with this microscope within one scan as well as between in consecutive scans to changes observed using scanning SAXS, demonstrating that the two sets of data are complementary. Thus, our approach enables quantitative analysis of radiation damage to cells in both single and consecutive scans *in situ* during scanning SAXS measurements.

## 2 Materials and Methods

### 2.1 Sample preparation

We employ human adrenal cortex carcinoma-derived SW-13 cells (ATCC CCL-105) (Leibovitz & McCombs III, 1973) that stably express fluorescent keratin hybrids (HK8-CFP, HK18-YFP) (Strnad *et al*., 2002; Windoffer *et al*., 2004). The cells are cultured in high-glucose (4.5 g L^*−*1^) Dulbecco’s Modified Eagle’s Medium (DMEM, D6429; Merck KGaA, Darmstadt, Germany) supplemented with 10 % (v/v) fetal bovine serum (FBS, 10270-106, Gibco), and 1 % (v/v) 100 U mL^*−*1^ penicillin and 0.1 g L^*−*1^ streptomycin (pen-strep, 15140122; Gibco). The cells are incubated at 37 ^*°*^C in a water-saturated atmosphere with 5% CO_2_. We culture the cells for one week after thawing and then add 50 g L^*−*1^ of geneticin (G418, L0015, Biowest, Nuaillé, France) to the medium as a selection agent.

Silicon nitride membranes (Si_3_N_4_, hereafter denoted by SiN, with a frame size of 5 mm *×* 5 mm, a frame thickness of 200 *µ*m, a window size of 1.5 *×* 1.5 mm, and a window thickness of 1 *µ*m; Silson Ltd, Warwickshire, UK) are plasma-activated at 50 W for 40 s (Zepto, Diener electronic GmbH & Co. KG, Ebhausen, Germany) to render the surface hydrophilic. Following the plasma activation, the membranes are immersed in ultrapure water and UV sterilized. Subsequently, the membranes are placed in 12-well cell culture plates, with the flat side oriented upward. To improve the cell adhesion, we apply 20 *µ*L of 5 % (v/v) fibronectin (F1141, Merck KGaA) in phosphate-buffered saline (PBS, D8537, Merck KGaA) to the membranes, then incubate them for one hour at 37^*°*^C. At about 80 % of confluence, the cells are detached from the culture flasks by adding 0.25 % trypsin with 0.02 % (w/v) EDTA (P10-020500, PAN-Biotech GmbH, Aidenbach, Germany) in PBS. The cells are then suspended in the culturing medium. The cells are then seeded onto the treated membranes at a concentration of 1.5 *×* 10^5^ cells mL^*−*1^ (6.25 *×* 10^5^ cells cm^*−*2^), and incubated for 24 hours. We chemically fix cells with 4 % methanol-free formaldehyde (28906, Thermo Fisher Scientific, Darmstadt, Germany) in PBS, and rinse them three times with PBS. The membranes with the fixed cells are individually stored in PBS with 1% (v/v) pen-strep and transported to the European Synchrotron (ESRF, Grenoble, France).

The resulting samples are referred to as ‘fixed-hydrated’. The fixation preserves native and higher-order protein structures in cells to a great extent. Storing the cells in a liquid environment, *i*.*e*., hydrated state, retains these preserved structures. Although we previously demonstrated that the fixation alters nanoscale cellular structures by comparing them with living cells (Weinhausen *et al*., 2014), they still preserve most structural information and are less sensitive to radiation than living cells (Weinhausen *et al*., 2014; Priebe *et al*., 2014), thus providing an excellent model system for our methods development.

### 2.2 Microfluidic sample environment

To maintain the cells’ close-to-physiological environment and in particular their hydrated state during the measurements, we use the microfluidic chamber as described in Ref. (Sinha *et al*., 2026), see Fig. 1d and e. Briefly, two SiN membranes, one hosting fixed-hydrated cells on its flat side and the other one being empty, are attached to the two sides of a home-built microfluidic device at the site of a punched hole as observation window. The device consists of two 8-*µ*m-thick polyimide foils (Kapton, SPEX SamplePrep, Metuchen, NJ, USA) that sandwich a channel structure made of Norland Optical Adhesive 81 (NOA81; Norland Products, Jamesburg, NJ, USA). The membranes are first attached to the hole in the device using thin double-sided tape (5 *µ*m thick; Nitto, Osaka, Japan) and sealed at their edges with cyanoacrylate adhesive (UHU, 478747-62, Bühl, Germany). This geometry confines the thickness of the liquid layer at the observation region to approximately 60 *±* 10 *µ*m. Thus, the background scattering is minimized while the cells remain in a hydrated state. Additionally, as the SiN is accessible by an objective, the chamber is compatible with optical microscopy. To connect the tubing for the inlet and outlet, as well as to provide mechanical stability to the chamber during the scanning, a ring-shaped polydimethylsiloxane (PDMS; Sylgard 184, Biesterfeld Spezialchemie GmbH, Hamburg, Germany) support is attached to the device. During the X-ray measurements and the microscopic imaging, degassed PBS flows through the channel at a rate of 100 *µ*L h^*−*1^, *i*.*e*., 0.132 mm s^*−*1^ in the observation region of the device. Therefore, the cells remain hydrated during the measurements.

**Figure 1:**
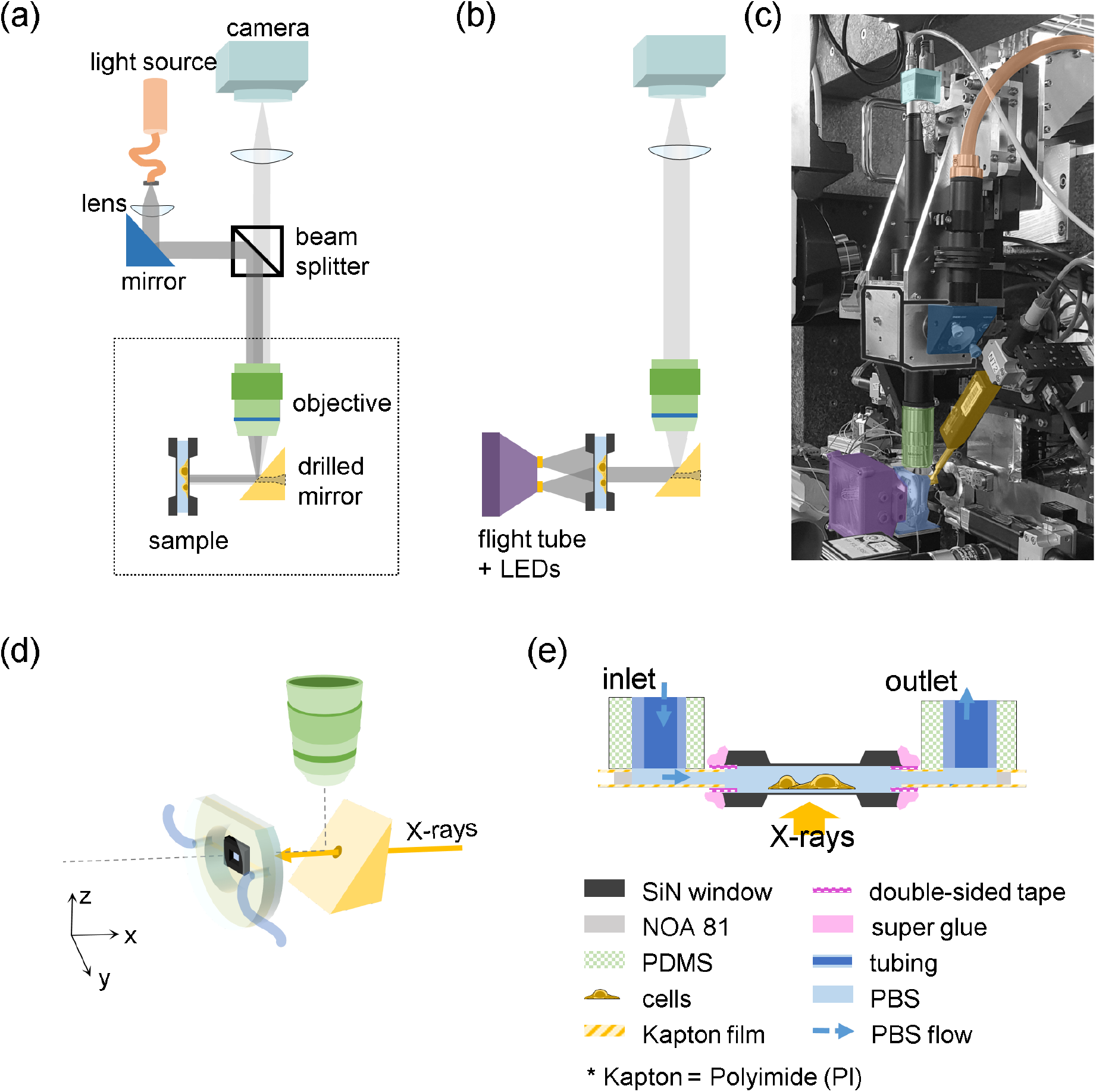
High-speed microscopy setup and dedicated sample chamber. (a) Schematic of the high-speed microscope in reflection mode and (b) in transmission mode (not to scale). For better visualization, all components are drawn in the same plane. Both modes share half of the optical path and components (from the sample to the camera); switching between the modes can be realized remotely by selecting the corresponding light source. (c) Photograph of the setup at beamline ID 13 at ESRF. The color masks indicate the optical components as shown in panels a and b. (d) Three-dimensional view of the chamber, prism mirror and objective for high-speed microscopy. The prism mirror reflects visible light from the sample to the objective, while the X-rays are guided to the sample through the drilled hole. (e) Schematic side view of the microfluidic sample chamber. Two SiN windows are separated by approximately 60 *µ*m inside a Kapton-NOA-Kapton microfluidic device.

### 2.3 Scanning small-angle X-ray scattering

We perform the combined scanning SAXS measurements and high-speed brightfield microscopy at the nano-branch of beamline ID 13 of ESRF. For the scanning SAXS experiments, the X-ray beam from the undulator is restricted to an energy of 15 keV using a Si-111 channel-cut monochromator. The beam is pre-focused by beryllium parabolic refractive lenses (Be-CRLs). The beam is further focused by a multilayer Laue lens with a working distance of 90 mm to a final size of 250 nm *×* 250 nm at a flux of 1.2 *×* 10^12^ photons s^*−*1^. At the endstation, the focused beam is conditioned by a 40 *µ*m *×* 40 *µ*m square-shaped PtIr aperture (order-sorting aperture, OSA) and enters the sample through a hole in the drilled prism mirror. The mirror is mounted on a separate motor stage for precise aliment with the X-rays. On the downstream side of the sample to reduce air scattering, we employ a helium-filled flight tube with an integrated LED array used for brightfield microscopy. The beamstop is located just outside exit window of the flight tube to block the direct beam. Scattered X-rays are recorded with a two-dimensional X-ray detector (Eiger X 4M; 2070 *×* 2167 pixels, *i*.*e*., *∼* 4 megapixels with a pixel size of 75 *µ*m *×* 75 *µ*m; Dectris, Baden, Switzerland). The sample-to-detector distance is about 1.9 m, as determined by measuring AgBeh powder in the empty chamber.

We employ the fast scan mode (Cassini *et al*., 2020; Yu *et al*., 2026). To track the radiation damage to the cells, we select multiple regions of interest (ROIs) with an area of 100 *µ*m *×* 100 *µ*m. Each ROI is scanned with a 0.5 *µ*m *×* 0.5 *µ*m step size and an exposure time of 5 ms; thus, a total of 40,000 scattering patterns are collected in approximately 4 minutes. We scan the same region 3 times with the same scan parameters to track the radiation damage by repeated scanning. Therefore, the time delay between two consecutive scans at the same measurement position is about 5 minutes (start time to start time), including overhead time. The average radiation dose *D* can be estimated by

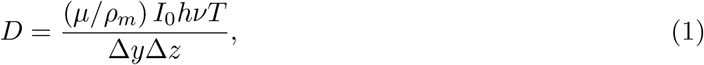

where *µ/ρ*_*m*_ = 1.60 cm^2^ g^*−*1^ is the ratio between the mass attenuation coefficient and the mass density of the cellular material (Berger *et al*., 2010), *I*_0_ is the photon flux of the incoming beam, *hν* is the photon energy, *T* is the exposure time per scan point, and Δ*y* and Δ*z* are the step sizes (Howells *et al*., 2009; Weinhausen *et al*., 2012). Thus, the radiation dose for the given scan parameters is 9.2 *×* 10^6^ Gy.

### 2.4 Data analysis

The scanning SAXS data are analyzed as described in Ref. (Yu *et al*., 2026). We determine the optimal *q* range as 0.19 - 0.39 nm^*−*1^ and use it to generate darkfield contrast images and to compute the anisotropy. A darkfield contrast image is obtained by calculating the total scattering intensity from the scattering center to the upper bound of the optimal *q* range, *i*.*e*., 0 to 0.39 nm^*−*1^. The anisotropy is defined by the circular variance of the azimuthal intensity profile within the optimal *q* range (0.19 to 0.39 nm^*−*1^) of the scattering patterns.

We use automated segmentation methods to determine the cellular regions and the background regions for background subtraction (Yu *et al*., 2026). In the darkfield contrast images, the background signal is defined as the average intensity of the pixels in the background region of each row. This signal is then subtracted from all pixels in the row to obtain the final darkfield contrast images. To compute the anisotropy, before computing the circular variance of each profile, we subtract the averaged signal corresponding to each row.

During the X-ray scans, we use our home-built high-speed microscope to record brightfield (BF) images of the scan region. These images are taken at the same frame rate as the X-ray scan (one image per scan point, *i*.*e*., 200 fps). We define the background signal as the average image of all BF images within a single dataset. To perform background correction, we normalize the BF images by the background image. While the sample continuously moves during the X-ray beam, the microscope images are taken, thereby recording a slightly differing field of view for each pixel. To account for these changes, we employ a FFT (fast Fourier transform)-based image registration method (Reddy & Chatterji, 1996). The displacement of the field of view in each image, relative to the initial image, is determined as the position of the maximum in the 2-dimensional cross-power spectrum between the two images.

To evaluate the BF images taken by the home-built microscope, we calculate the contrast-to-noise ratio (CNR) of 10 cells in an example image. First, we manually generate cell masks and calculate the centroid of each cell mask. Then, we define a square region centered on the centroid with a size of 226 *×* 226 pixel^2^, or 108 *×* 108 *µ*m^2^. This region restricts the selection area to the vicinity of the target cell when setting the reference background intensity distribution for further steps. If the selected region exceeds the image boundary, the excess area is disregarded. Thus, each area contains at least one cell. Next, we manually select a rectangular background area in each square to calculate the background reference value. The CNR of each cell is defined by

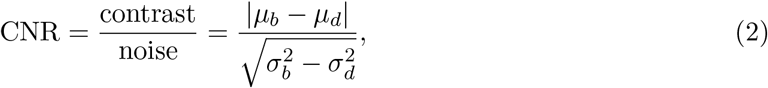

where *µ* is the mean and *σ* is the standard deviation of the selected bright (*b*) and dark (*d*) areas (Timischl, 2015). Each pixel on the cell mask falls into one of three categories: bright, dark, or background. The bright area is defined as the intensity being higher than the 99th percentile of the reference background intensity distribution. Similarly, the dark area is defined as the intensity being lower than the 1st percentile of the distribution. The remaining pixels are considered part of the background. The CNR is then calculated from the categorized pixel subsets within the cell mask.

We use gradient magnitude maps to quantitatively track changes in the BF image during a single scan or between consecutive scans. We use the Sobel operator to calculate the image gradient (Khan *et al*., 2019), a two-component vector (*g*_*y*_, *g*_*z*_), combining the derivatives of intensity in each point in the original image along the *y−* and *z−*axes. The gradient magnitude is then defined as the length of the vector, *i*.*e*., 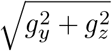. Although the Sobel operator introduces an slight orientation bias, it does not affect the magnitude.

## 3 Results and discussion

### 3.1 *In situ* high speed microscopy

Imaging biological samples by X-rays is highly attractive for several reasons, including the intrinsic high resolution, high penetration power, the ability to image label free specimens, and the combination of information gathered in real and reciprocal space, *i*.*e*., images and structural information. However, one great challenge remains the extreme sensitivity of biological matter to radiation and the changes in the sample associated with this sensitivity. Samples in aqueous environment are even more prone to radiation damage than dry samples, but closer to biological reality and therefore of high interest. To track radiation damage in hydrated biological cells during X-ray measurements, an *operando* imaging approach is needed. We build a high-speed microscope and integrate it into the scanning SAXS setup at the nanobranch of the beamline ID 13 at the ESRF. One key component for the *in situ* imaging is a prism mirror with a hole located in the X-ray beam path, as shown in Fig. 1d. Specifically, we employ a right-angle prism mirror with 10 mm edges (MRA10-P01, protected silver; Thorlabs GmbH, Newton, NJ, USA). We drill a hole of approximately 0.4 mm diameter through the center of the inclined side towards one flat side; we then drill a hole of 3 mm diameter and 1.5 mm depth from the flat side. One of the prism’s sharp edges is grounded down so that the length of the flat side without the hole is 6 mm. This way, the prism mirror can be moved closer to the sample, which is an important factor for the tight setup at the beamline. See Fig. S1 in the supporting information for details on the prism mirror.

This microscope can be operated in two imaging modes, reflection and transmission, see Fig. 1a and b, respectively. For reflection microscopy, we use a white high-power LED light source (orange; intraLED 5, Fiber Optic, Spreitenbach, Switzerland). First, light from the source enters a collecting lens (plano-convex lens, LA1805-A, diameter of 25.4 mm, focal length 30 mm; Thorlabs). Then, the light reflects off a mirror (blue; BBE1-E02, broadband dielectric-coated elliptical mirror; Thorlabs), which changes the propagation direction of the light. Next, it enters a beam splitter (black; BSW16, diameter of 2 inch, thickness 8 mm, coating for 400 nm to 700 nm at a 45^*°*^ incidence angle, a 50:50 split ratio; Thorlabs). After passing through the beam splitter, the light illuminates the sample *via* an objective (green; Plan Apo, NA 0.28, working distance 34 mm; Mitutoyo Deutschland GmbH, Neuss, Germany) and the drilled prism mirror (yellow). We employ a long working distance objective, which provides sufficient distance between the objective, the drilled prism mirror, and the sample (see Fig. 1d). The light reflected from the sample follows the same optical path back to the beam splitter and is then focused on the camera by a tube lens (TTL200-A, working distance of 151.8 mm; Thorlabs). This optical path is illustrated in Fig. 1a. Transmission microscopy follows the same optical path from the sample to the camera (light blue; acA640-750um, frame rate 751 fps, pixel size of 4.8 *µ*m *×* 4.8 *µ*m, sensor size of 3.07 mm *×* 2.3 mm; Basler AG, Ahrensburg, Germany).

To transform this setup into a transmission microscope, an light source on the side of the sample opposite of the objective is required. We employ a custom-made, square-shaped frame with a 12-LED array (20 mA intensity each). The LED array is mounted on the upstream window of the flight tube. A photograph of the final setup is shown in Fig. 1c. Since both microscopes share most of the optical path, the two modes can be easily switched by turning the corresponding light source on or off. This versatility makes the microscope ideal for imaging both opaque samples, such as the X-ray resolution target (“Siemens star”), and transparent samples, such as the hydrated biological cells.

From the software side, we fully integrate this microscope into two control systems: the beamline control system and the web-based graphical interface control system, Daiquiri (Fisher *et al*., 2021). This integration enables remote control of the microscope (imaging modes, illumination intensity) and management of data acquisition. We employ a high-speed camera to acquire images at the same rate as the X-ray scan, *i*.*e*., one full microscopy image per scan point. The X-ray detector triggers the imaging acquisition, thereby synchronizing the X-ray data and imaging acquisition. Consequently, the microscope camera has a minimum acquisition time of 2 ms, including overhead time, allowing for high-speed and real-time imaging.

Fig. 2a shows an example microscopy image in transmission mode of fixed-hydrated cells in the microfluidic chamber taken with the high-speed microscope. This image is the first image of an entire series, recorded during a SAXS scan. The SAXS scan is performed on an area of 100 *×* 100 *µ*m^2^ (black box in Fig. 2a, with a step size of 0.5 *µ*m in both directions and an exposure time of 5 ms at each position. Thus, the sample is moved 200 steps horizontally and vertically each, and a total of 40,000 SAXS patterns and microscope images are collected. Despite the short exposure time, the outlines of the cells are distinguishable from the background (see also Fig. S2a,b in the supporting information).

**Figure 2:**
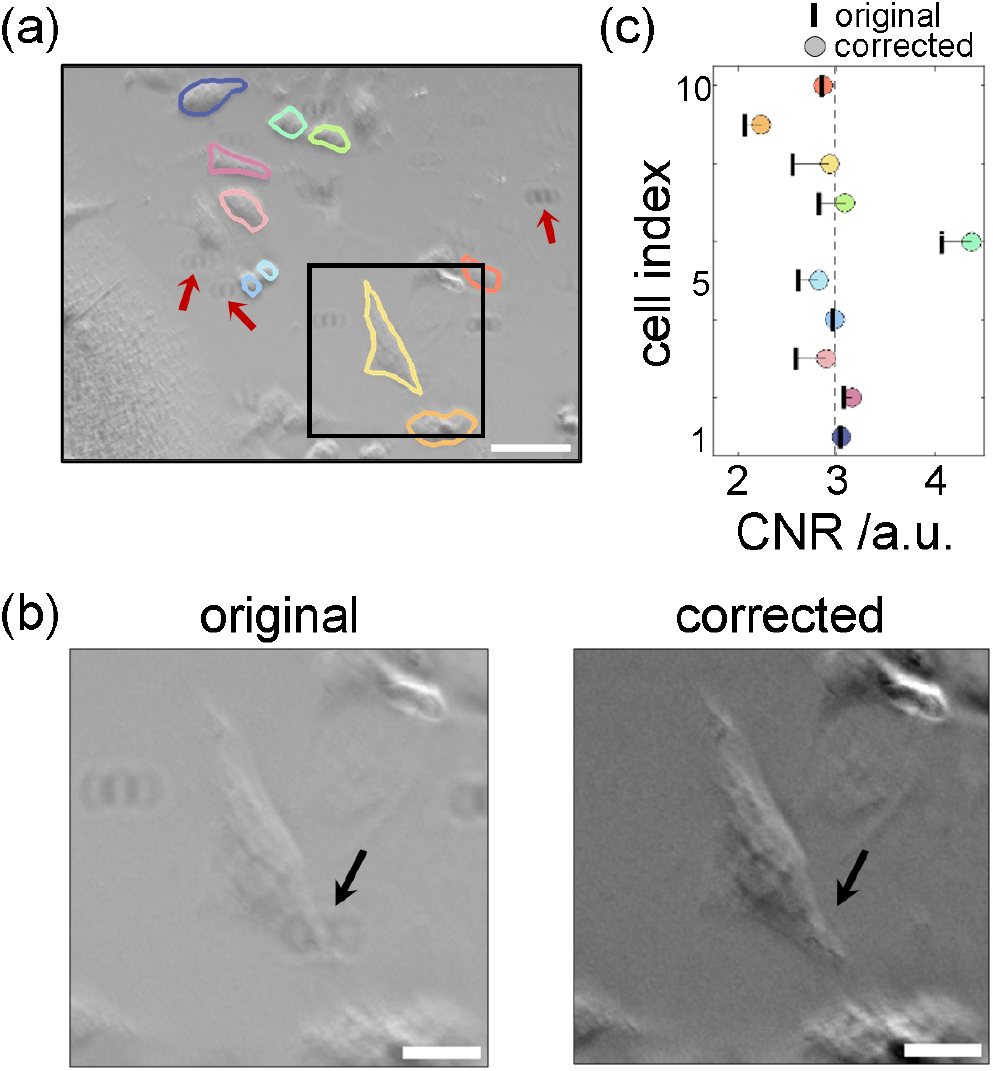
(a) Example brightfield image taken with the high-speed microscope. 10 cells are labeled by manual segmentation. Red arrows: ring-shaped patterns. (b) Zoomed brightfield images of the selected region (black box) in panel a; (left) original image; (right) after the background correction. Arrows: one example of a ring-shaped pattern which is removed after the correction. Scale bar in panel a: 50 *µ*m; in panel b: 20 *µ*m. (c) Contrast-to-noise ratio comparison before (original) and after (corrected) the background correction of 10 regions of interest (226 pixels *×* 226 pixels), each containing at least one of the cells labeled in panel a (see color code). The dashed line indicates a CNR of 3.

The images recorded with the high-speed microscope show characteristics similar to those of oblique microscopy. One such characteristic is that the cells appear as pseudo-3D with directional shadows, similar to what is known from differential interference contrast (DIC) microscopy. This makes the oblique microscope useful for imaging transparent specimens, proving better contrast and resolution than coaxial illumination (Sanchez *et al*., 2018). Thus, this illumination enables us to observe the cell shapes despite the short exposure times. Nevertheless, the image quality is not optimal, see Fig. 2b, left. Additionally, the illumination generates an inhomogeneous background intensity in the sample plane (see Fig. S2c,d in the supporting information) and we observe ring-shaped patterns presumably produced by the microscope optics (see the red arrows in Fig. 2a). These artifacts affect the image quality and alter the background signal of the same cells measured in different positions during scanning. Thus, they must be corrected for further quantitative analysis. We perform a flat-field correction to address the artifacts. Artifacts arising from non-uniform illumination and spatial variations in the optical response can be modeled as multiplicative noise. Therefore, the collected image, *I*_*col*_, can be described by

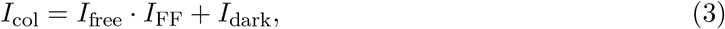

where *I*_free_ is the artifact-free image, *I*_FF_ is the flat-field image including artifacts, and *I*_dark_ is an image taken with the shutter closed and contains intensities originating from the camera’s readout noise and dark current (Pang *et al*., 2012). The flat-field image contains global and local artifact information. However, this local information is negligible when analyzing the entire set of images; therefore, *I*_FF_ can be estimated from only the global artifact information. Furthermore, we can neglect the dark image *I*_dark_, because the dark pixel intensities are very small at a short exposure time (Pang *et al*., 2012). Therefore, *I*_free_ can be simply estimated by dividing *I*_col_ by *I*_FF_. We calculate *I*_FF_, *i*.*e*., an average microscope image of all individual recordings collected during a single scan (see Fig. S2c in the supporting information for an example). In each microscope image, the position of the observed object, *i*.*e*., cells, continuously changes, while the position of the artifacts remains constant. Therefore, the common features present in all images remain after the averaging, while the position-dependent features disappear. To avoid outliers, we calculate the truncated mean at each pixel by discarding values smaller than the 1^st^ quartile or larger than the 3^rd^ quartile. Then, we divide *I*_*col*_ by *I*_FF_. This correction removes the ring-shaped artifacts, see the black arrows in Fig. 2b, left *versus* right, and calibrates the background illumination, see Fig. S3 in the supporting information. Additionally, the flat-field correction improves the overall contrast between the cell and the background, and the cell contours in the corrected image are clearer than in the original image, because the correction removes the global shading, making the true contrast visible.

We quantify the improvement in image quality by calculating the CNR, as described in Eq. 2. The CNR is independent of the image intensity scale and offset, both of which are affected by the given illumination intensity and exposure time (Burgess, 1999; Timischl, 2015). Therefore, the CNR is suitable for evaluating the performance of a microscope with specific types of samples. We manually select 10 cells from the raw image across the entire field of view, including the SAXS-scanned cell (the cell in the black box in Fig. 2a). The selected cells are annotated with different colors in Fig. 2a, and their CNR values are shown with the same color annotation in Fig. 2c. As shown in Fig. 2c, the CNR values all increase in the corrected version (colored circles) compared to the original (black bars). Despite the varying extent of the increment, the overall improvement demonstrates that the flat-field correction enhances the image quality.

After the correction, most CNR values are close to 3 (see dashed line in Fig. 2c), which can be interpreted using the detection limit (Kaiser, 1965) and the Rose criterion (Rose, 1948). The detection limit is typically defined as the three times the standard deviation of the background signal (3*σ*), and is the minimum threshold at which a signal collected by a detector can be distinguished from noise. Thus, the detection limit describes an instrument’s performance. The Rose criterion refers to the visibility of a feature in an image to the human eye and brain. Note that we applied the Rose criterion to the CNR, not to the signal-to-noise ratio (SNR), because the CNR provides a more precise estimation when the signal is very weak (Bao & Chatziioannou, 2010; Dieckens *et al*., 2013). Taken together, the CNR values close to 3 indicate that the signal is at a reliable level, but insufficient to be distinguished by the human eye. This implies that conventional image analysis techniques, which require human involvement, cannot be used to analyze these images and a different approach is needed.

### 3.2 *Operando* tracking of radiation damage

Fig. 3a shows an example series of microscope images recorded during a single SAXS scan. For the analysis, we crop the microscopy images to the scanned area. Here, we show the first image of selected scan rows, based on the features of the scanned cell (see black outlines): (i) row 1, the starting point of the scan (see also Fig. 2b, right), (ii) row 80, the first row on the cell nucleus, (iii) row 120, the last row of the cell nucleus, (iv) row 160, the last row on the cell, (v) and row 200, the last row of the scan. The white dashed lines denote the corresponding scan positions in each image. All images are flat-field corrected.

**Figure 3:**
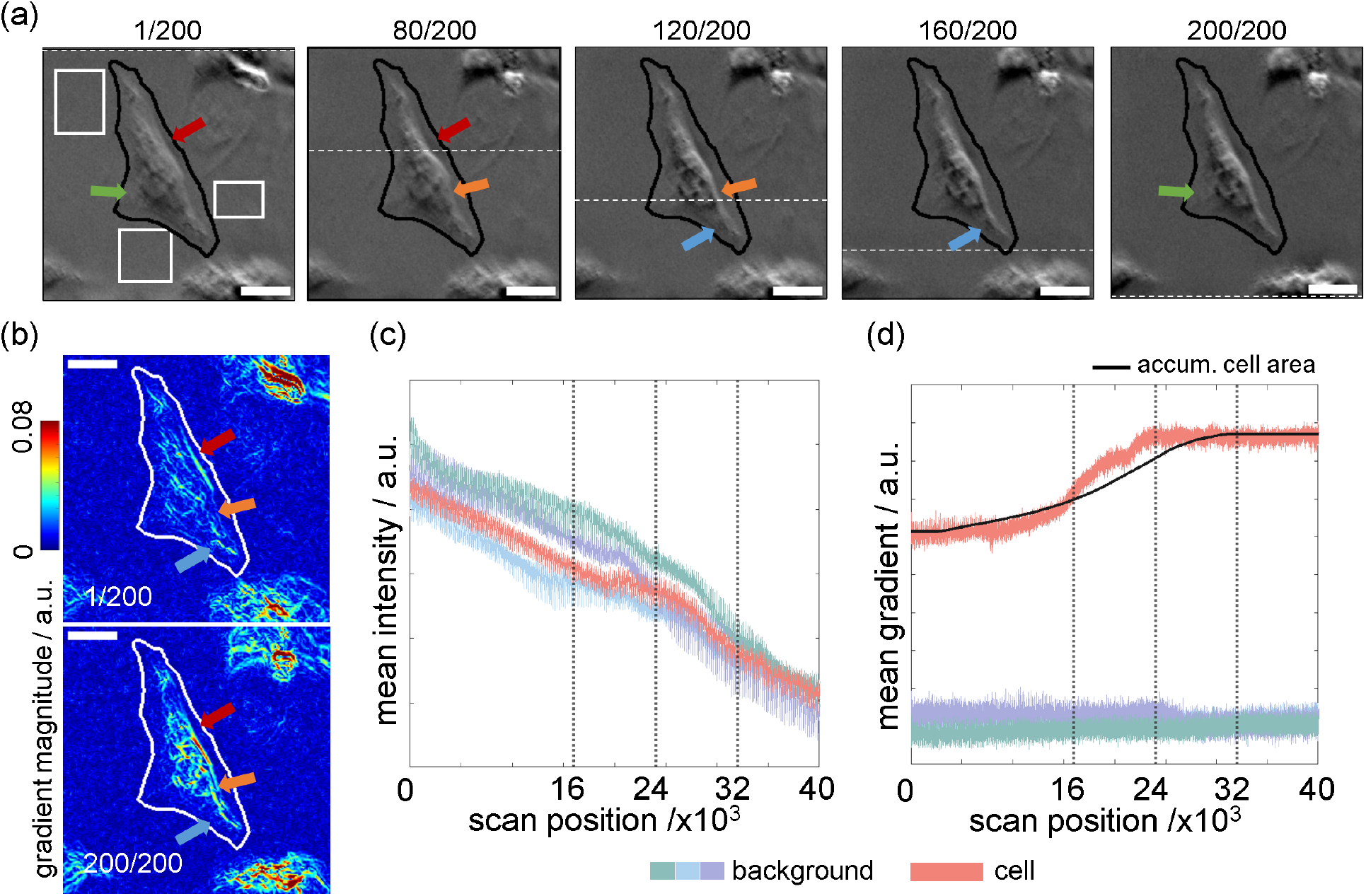
Radiation damage tracking during scanning SAXS. (a) Series of brightfield images taken during a single X-ray scan. Among 40,000 images within a single scan (*i*.*e*., one per scan position), the first images of the first, last, and 3 intermediate rows are selected. The row numbers are provided at the top. Four regions are manually selected to compare the intensity and mean gradient changes by scan position, *i*.*e*., three background regions (white boxes) and a cell region (black outline). The colored arrows indicate positions where changes are visible within the presented image series. (b) Example gradient maps of the first and the last image of panel a. The gradient map is calculated by the Sobel operator and plotted on a color scale. (c) Intensity and (d) mean gradient changes of the four selected regions in panel a; red shows the cell region. The black dashed lines indicate the frames shown in panel a. The black solid line in (d) shows the accumulation plot of the cell mask area obtained from the darkfield contrast image (scanning SAXS).

As the scan progresses (from left to right), changes appear in the cell features. We mark some of these features with colored arrows in the individual images. The colored arrows are in the same position in each image. The first notable change is that the cell shrinks in size. The bright cell contour marked by the red arrow in row 80 is further away from the cell mask boundary (black solid line) than the one in row 1. This change is also visible in the orange-marked region when comparing row 80 to row 120.

The second change is that the contrast within the cell region is locally enhanced. Regions that are brighter than their surroundings in row 1 become even brighter (see the red arrows), and regions that are darker than their surroundings become darker (see the blue arrows). Additionally, the darker region marked by the orange arrow in row 80 shifts upward in row 120 and becomes darker. This result is a combination of cell shrinking and contrast enhancement.

The observed changes occur in the direction of the scan (from the top to the bottom of the image). As the images taken at row 80 and row 120 show, these changes are only present in the upper part above the dashed white line. This demonstrates that the X-ray radiation locally induces these changes. Nevertheless, some parts of the cell contours remain blurred before and after the exposure (see the green arrow in row 200). Therefore, an adapted approach to tracking these changes is necessary. To quantify the changes observed in the microscopy images, we analyze gradient magnitude maps. To this end, we apply the Sobel operator to a total of 40,000 images, as it provides high computational speed and reliable results. The gradient map indicates high values where the local contrast is high. Fig. 3b shows examples of two gradient maps computed from the first and last images shown in panel a.

Comparing the maps of row 1 and 200, we see our observations from the brightfield images confirmed. The bright, thick line (red arrows) shifts toward the cell center, consistent with Fig. 3a. Additionally, the map from row 200 shows thicker, more intense features than the one from row 1. Short lines on the map from row 1 get extended and even merged and increase in magnitude (orange and blue arrows). Since the gradient magnitude represents the local contrast information, these changes represent the local contrast enhancement in the microscope images due to X-radiation.

The radiation damage is effective on the cell area (black outlines in Fig. 3a) but not the background (white boxes). However, when comparing the mean intensities Fig. (3c) for these four regions plotted against scan position, they cannot be distinguished. Although the intensity changes due to the X-ray exposure render the bright areas brighter and the dark areas darker, they mutually compensate when calculating the mean, despite enhancing the image contrast. All four curves decay continuously as the scan progresses. The decay is presumably due to, first, “shadowing” of the chamber holder that removes some of the transmitted light and, second, X-radiation-induced changes. This assumption is confirmed by the fact that when rescanning the same region several times, the mean intensity starts at a high value for each scan, albeit not as high as the first one, and then decreases during the scan.

A better measure to distinguish cells from background is the mean gradient, see Fig. 3d. All values in the cell curve (red) are considerably higher than those in the background curves (blue, green, purple). The curve also follows a smooth step function, starting with a gradual increase, followed by a sharp rise, and then a saturation. To interpret the shape of the curve in relation to the shape of the cell, we added black dashed lines corresponding to the selected scan lines in Fig. 3a. The gradual increase in the first region (from row 1 to 80) results from the X-ray exposure to the upper cytoplasmic region. The sharp rise between lines 80 and 120 represents the radiation damage to the nuclear region. From row 120 on, the curve is saturated. This is probably because the cell area in this range is small, thus contributing only little. The increasing trend is consistent with the two findings above, *i*.*e*., the contrast enhancement and the changes occurring along the scan direction. We also compare this curve with the accumulated cell area curve obtained from the scanning SAXS measurement (black solid line). This curve represents the total number of scanned pixels from the cell region in the darkfield contrast image, plotted against the scan position (see section 2 for details). Both curves demonstrate a very similar trend, showing that the curve shape is related to the cell shape. Taken together, we can quantify the changes observed in the microscope images induced by X-ray exposure by the gradient magnitude. An additional data set is shown in Fig. S4 in the supporting information, see also movie S1.

### 3.3 Simultaneous scanning SAXS and light microscopy

To track the radiation damage caused by repeated X-ray exposure, we scan the same region three times (scans 1 to 3) with the same scan parameters, *i*.*e*., a step size of 0.5 *µ*m in both directions and an exposure time of 5 ms at each position. Including the overhead time at the beginning of each scan and between the scan lines, each scan position is exposed to X-rays every 5 minutes. The high-speed microscope records images during the three scans in synchrony with the X-ray exposures. We use the first microscope image from each scan for damage tracking. The image from scan 1 captures the intact cell, while the images of scans 2 and 3 contain the image of the cell after the first and second exposures, respectively.

Fig. 4a shows three gradient magnitude maps from each scan. The first image (scan 1) is the same as in Fig. 3b. The changes from scan 1 to scan 2 are consistent with the changes observed from the row 1 to row 200 maps in Fig. 3b. However, there are only subtle changes in line shapes and magnitude from scan 2 to scan 3. This indicates that the first exposure significantly alters the local contrast of the cell region in the microscope image, whereas additional exposures have little effect. Fig. 4b shows the darkfield contrast images obtained from the SAXS scans 1 to 3. The shape of the cell is clearly distinct from the background in all the images, in contrast to the microscope images in Fig. 3a. The cell body region and the nucleus region can also be distinguished by variations in intensity. These intensity variations result from non-uniform mass distributions within the cell (Weinhausen *et al*., 2012). These characteristics suggest that the quality of the collected SAXS data is sufficient for a quantitative analysis to be performed on the cell and nucleus regions. We overlay the cell contour (white) and nuclear contour (black), obtained from the darkfield contrast image from scan 1 using the automatic segmentation method described in Sec. 2.4, onto all images in the series to enable further visual comparison and numerical analysis.

**Figure 4:**
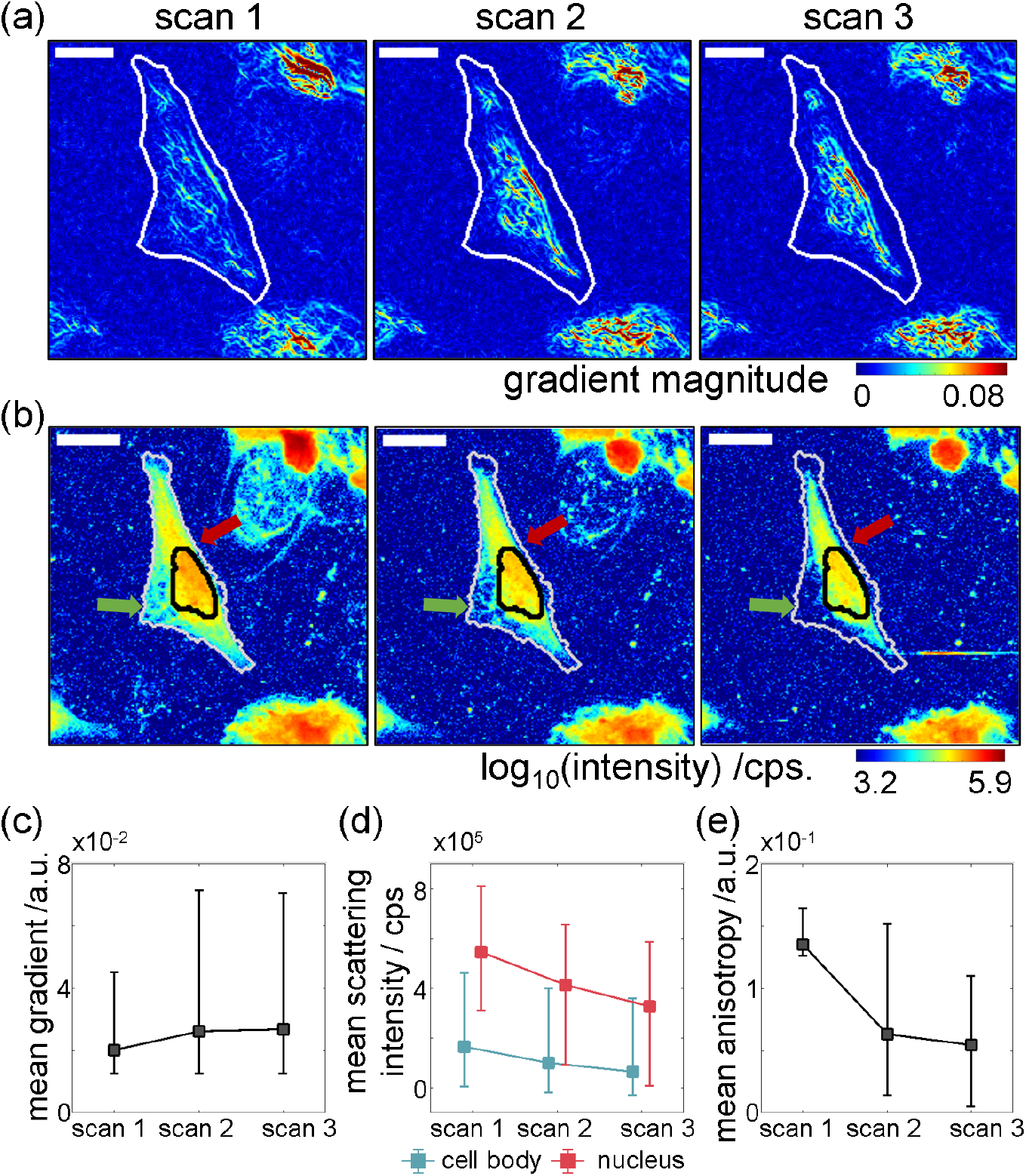
Multiple re-scanning of the same region and associated radiation damage effects. (a) Gradient maps calculated from the first brightfield image of each scan. (b) X-ray darkfield contrast images. Arrows indicate significant changes between scans. Scale bars in panel a and b: 50 *µ*m. (c) Mean gradient magnitudes within the cell region (white outline in a), plotted against scan number. (d) Mean scattering intensity plotted against scan number for the nucleus region (red, see black outline in panel b) and the cell body (blue, see white outline in panel b). (e) Mean anisotropy of the selected pixels, *i*.*e*., higher than the 99th percentile of the background region within the cell body region plotted against scan number. The error bars in panel c to e denote the first and the 99th percentiles of the distributions.

In the darkfield contrast images, radiation damage alters both the cell contour and the scattering intensity of the pixels within its region. In scan 2, the cell contour shifts inward compared to that of the first scan. Where the area of the cell shrinking is well visible in the gradient maps in Fig. 3b, this change is clearly observed in the darkfield images (red arrows). Also, the scattering intensity of the pixels within the cell region decreases. The nucleus region (black outline) turns from red to yellow, and the cell body region changes from yellow to cyan or from cyan to dark blue. Notably, some pixels in the area where the cell contour is blurred in the microscope image in Fig. 3a have a similar intensity as the background region (green arrows). This indicates that the cellular material at this position is lost due to the radiation damage.

Comparing the darkfield images from scan 2 and 3, the changes indicating radiation damage proceed. The cell contour shifts further inward compared to the scan 2 image (red arrow), and the overall intensity further decreases. The pattern of continuous changes from scans 1 to 2 and 2 to 3 is clearly distinctive compare to what we observed in the gradient maps.

After this qualitative assessment of the gradient maps derived from microscopy and the darkfield images derived from scanning SAXS, we turn to a more quantitative analysis. We compare the changes from both image series in Fig. 4 panels a and b. To this end, we first compare the mean gradient magnitude of the cell region (white outlines in Fig. 4 a) against scan numbers in Fig. 4c. The error bars correspond to the first and 99th percentile of all pixel magnitudes. The mean value increase from scan 1 to 2, as does the error bar, and stays constant from scan 2 to 3, in agreement with our qualitative analysis of the gradient magnitude maps in Fig. 4a. This finding suggests that the local contrast in the cell region primarily changes during the first exposure and that the subsequent exposure does not yield significant changes.

By contrast, the mean scattering intensity (Fig. 4d) shows a continuous downward trend. We plot the mean scattering intensity of the nucleus region (the black-outlined region, red) and the cell body region (the region between the black and white outlines, blue). Notably, the nucleus region has higher overall mean intensities than the cell body region because the nucleus has a higher mass density and it is thicker, thus providing more material in the beam path. The continuous changes in the mean scattering intensity in both regions suggest that mass loss in the cell occurs continuously through repeated exposure.

We additionally analyze the anisotropy of the scattering patterns to track damage to the ordered structures within a cell, as described in Ref. (Yu *et al*., 2026). To this end, we select pixels in the cell body region with anisotropy values above the 99th percentile of the background regions in scan 1 and track their changes. Fig. 4e shows the mean anisotropies of the selected pixels from scans 1 to 3. The mean anisotropy decreases considerably from scan 1 to 2, and much less from scan 2 to 3. This suggests that the first exposure damages most of the ordered structures within the cell. The subtle decay from scan 2 to 3 indicates the structures remaining after the first exposure are further damaged, as discussed in Ref. (Yu *et al*., 2026). Taken together, we can quantitatively track radiation damage to cells by combining two image modalities, high-speed microscopy and scanning SAXS in an *operando* manner. Analyzing the obtained images with different analysis approaches reveals various cellular characteristics. Therefore, this approach allows us to investigate how cells respond to repeated X-ray radiation in a complementary and directly correlative manner.

## 4 Summary and conclusion

We present a high-speed, on-axis brightfield microscope and quantitatively track radiation-induced changes in single biological cells during a SAXS scan. The microscope combines an on-axis configuration for *in situ* imaging with a high-speed camera to achieve rapid acquisition rates. This enables synchronous imaging while recording scattering patterns. The recorded images show local contrast changes within the cell area as the scan progresses, indicating that X-rays alter cell morphology. The cell features in the micrographs show insufficient contrast for conventional image analysis approaches. However, we demonstrate that these changes can be tracked quantitatively using image gradient magnitude maps, which show good agreement with the observed changes in the micrographs. Interestingly, when applying our approach to repeated scans of the same cell region, we observe significant changes during the initial exposure, whereas subsequent exposures result in only minor changes. The changes detected from the bightfield microscopy images detected by the gradient magnitude maps are similar to those observed in mean anisotropy in scanning SAXS, but differ from the continuous decrease in the mean scattering intensity. Therefore, this multimodal imaging technique provides complementary information by the different methods. Nevertheless, we are currently limited by the image quality and future improvements will even increase the applicability of the approach. Our study demonstrates that radiation damage to biological cells can be tracked and analyzed quantitatively using a high-speed, *in situ* imaging technique during single and repetitive SAXS scans. Thus, the method can be used to optimize the measurement parameters further, resulting in good signal quality with minimized X-radiation damage. This multimodal approach provides complementary information, opening up the potential for multifaceted investigations of cellular responses to X-ray radiation. This approach can also be applied to other cell types or to living cells in a straightforward manner.

## Acknowledgements

We are grateful for the beamtimes associated with the long term project SC-5436 granted by ESRF (The European Synchrotron, Grenoble, France), and in particular outstanding support by the ID 13 staff and the PSCM (Partnership for Soft Condensed Matter), specifically Diego Pontoni, François Villar, Michael Hahn, Delphine Baboullin, Bastien Harnoux, Daniel Fiole, Pierre Lloria, and Peter van der Linden. We also thank Peter Luley for technical support, Peter Gawlitza and Sven Niese for providing MLL-lens prototypes used for these experiments.

## Data availability

Upon acceptance of the manuscript, the data will be uploaded to a repository (GRO.data) and the respective doi will be included here.

