## Supporting information for "In situ radiation damage tracking during scanning small-angle X-ray scattering"

### 1 Design of the drilled prism mirror

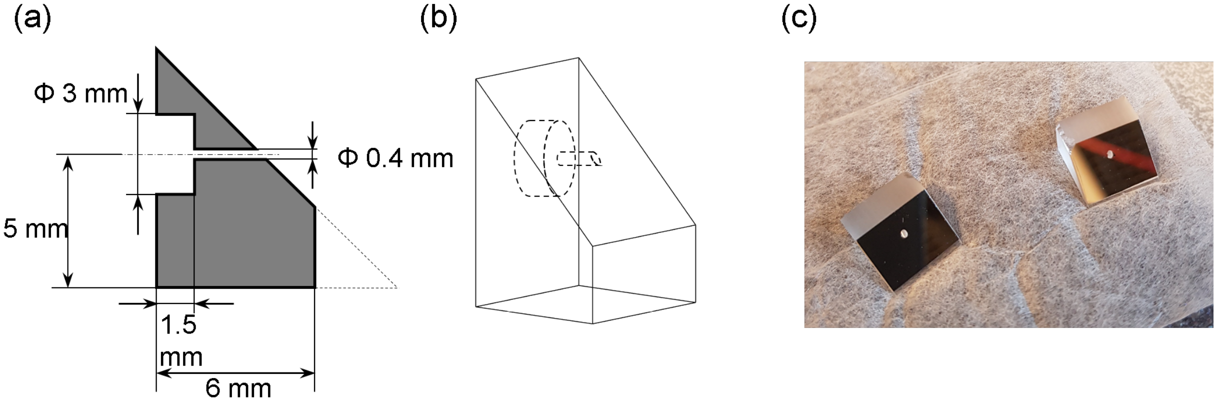


Figure S1: (a) Cross-sectional view of the drilled prism mirror. (b) 3D projection of the design. (c) Photograph of the mirrors after drilling.

#### 2 The effect of the oblique illumination

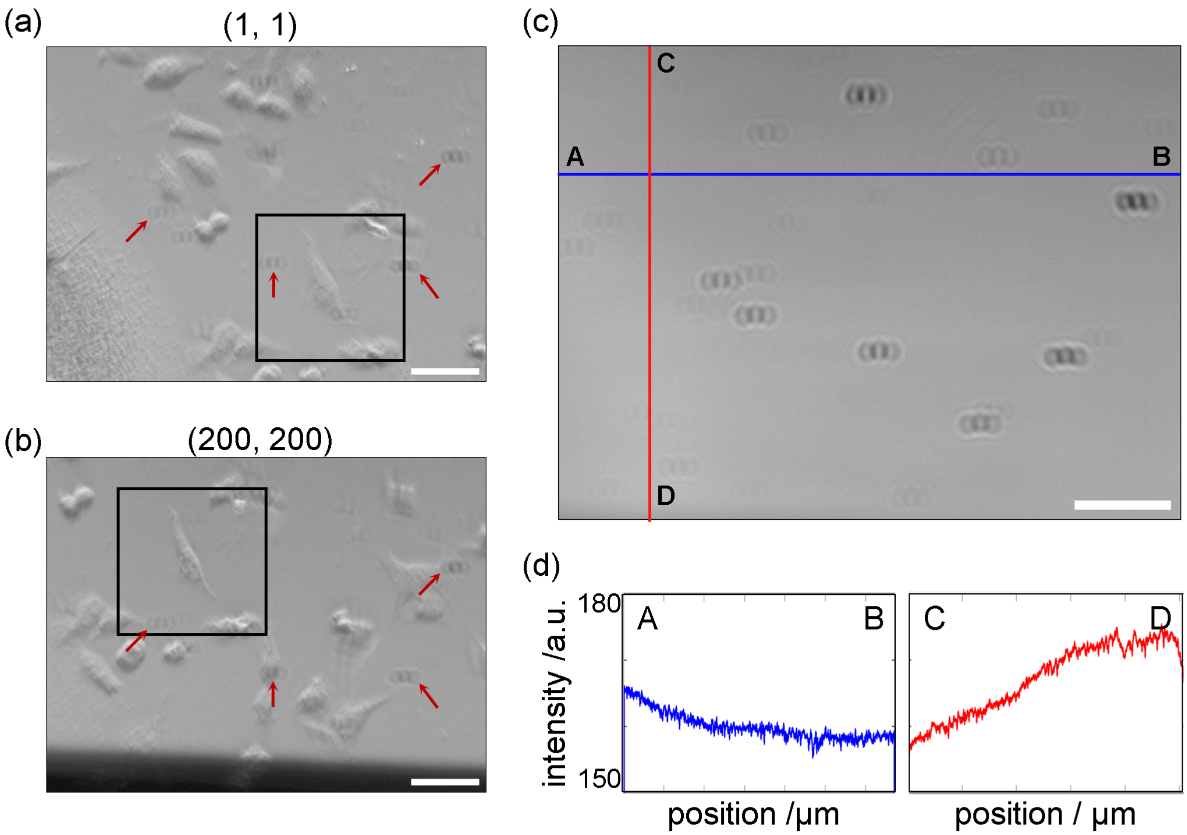


Figure S2: The effect of the oblique illumination. Raw microscope images of (a) the first and (b) last position of the SAXS scan. The index is read as (column, row). Panel a is the same image as shown in Fig. 2a in the main text, but without cell outlines. The cells appear pseudo three dimensional. The black boxes mark the SAXS scan region for this scan. Although the relative position of the cells changes throughout the scan, the position of the artifacts remains the same. (c) Truncated mean image of a single dataset (scan), which we use as flat-field image for image correction. (d) Two line profiles across the lines A to B and C to D in panel c, showing the intensity gradient across the field of view. Scale bars: 50 $\mu$m.

#### 3 Flat-field correction

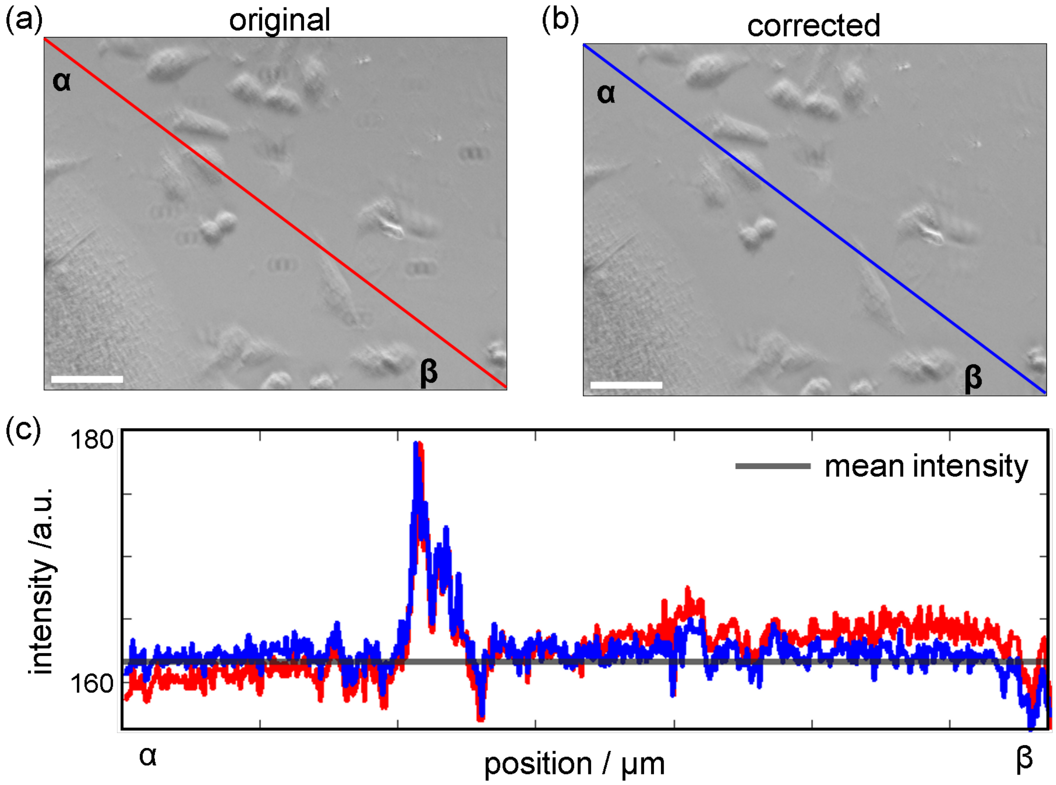


Figure S3: Comparison of an example microscopy image before and after the flat-field correction. (a) Original and (b) corrected microscopy image. (c) Two line profiles across the line from $\alpha$ to $\beta$. The red line stems from the original image, the blue line stems the corrected image. The black solid line shows the average intensity of the corrected image. The red line gradually increases from $\alpha$ to $\beta$, indicating the uneven background illumination. By contrast, most of the intensities in the blue line are close to the mean intensity, indicating the background illumination is flat after the correction. Scale bars: 50 $\mu$m.

#### 4 Additional dataset: radiation damage tracking during scanning SAXS

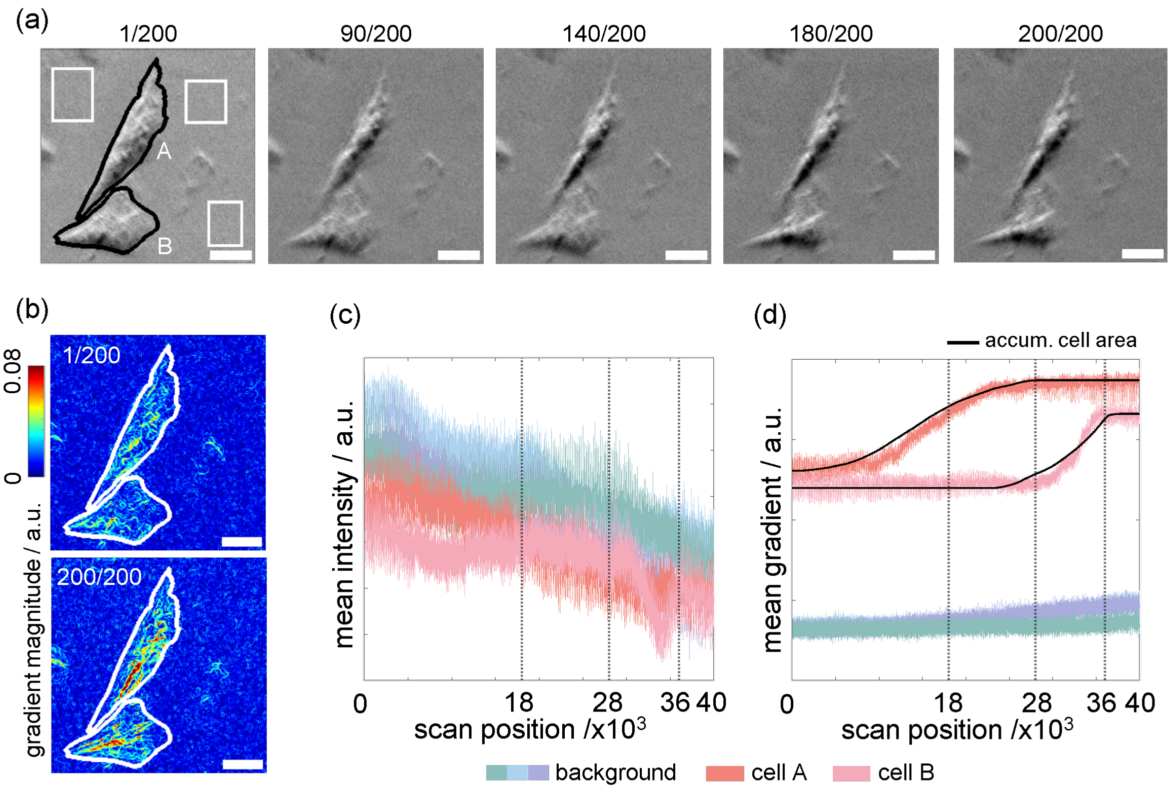


Figure S4: Radiation damage tracking during scanning SAXS (see Fig. 3 in the main text) of an additional dataset. (a) Series of bright-field images taken during a single X-ray scan. Among 40,000 images within a single scan (i.e., one per scan position), the first images of the first, last, and 3 intermediate rows are selected according to the cell shape. The row numbers are provided at the top. Five regions are manually selected to compare the intensity and mean gradient changes by scan position, i.e., three background regions (white) and two cell regions (black). Two cells are marked by A and B. (b) Example gradient maps of the first and the last image of panel a. The gradient map is calculated by the Sobel operator and plotted on a color scale. (c) Intensity and (d) mean gradient changes of the five selected regions in panel a; red and pink indicate the cells A and B, respectively. The black dashed lines indicate the frames shown in panel a. The black solid lines in d show the accumulation plot of each cell mask area obtained from the dark-field contrast image by scanning SAXS.

### 5 In situ imaging of biological cells during scanning SAXS

Movie S1: In situ imaging of biological cells during scanning SAXS measurement: (Left) Time-lapse brightfield video recorded during scanning; (Center) Corresponding gradient maps; (Right) Darkfield contrast image generated from the scanning SAXS data of the same dataset in as shown in Fig. 3 in the main text. The first image of each scan line is used to generate the video. The time needed to scan one line is approximately 1.3 s and the video is accelerated to fps. Scale bar: 20 $\mu$m.
